# EpiAlignment: alignment with both DNA sequence and epigenomic data

**DOI:** 10.1101/562199

**Authors:** Jia Lu, Xiaoyi Cao, Sheng Zhong

## Abstract

Comparative epigenomics, by subjecting both epigenome and genome to interspecies comparison, has become a powerful approach to reveal regulatory features of the genome. Thus elucidated regulatory features surpassed the information derived from comparison of genomic sequences alone. Here, we present EpiAlignment, a web-based tool to align genomic regions with both DNA sequence and epigenomic data. EpiAlignment takes DNA sequence and epigenomic profiles derived by ChIP-seq, DNase-seq, or ATAC-seq from two species as input data, and outputs the best semi-global alignments. These alignments are based on EpiAlignment scores, computed by a dynamic programming algorithm that accounts for both sequence alignment and epigenome similarity. For timely response, the EpiAlignment web server automatically initiates up to 140 computing threads depending on the size of user input data. For users’ convenience, we have pre-compiled the comparable human and mouse epigenome datasets in matched cell types and tissues from the Roadmap Epigenomics and ENCODE consortia. Users can either upload their own data or select pre-compiled datasets as inputs for EpiAlignment analyses. Results are presented in graphical and tabular formats where the entries can be interactively expanded to visualize additional features of these aligned regions. EpiAlignment is available at https://epialign.ucsd.edu/.

## INTRODUCTION

Epigenomes play pivotal roles in cell identity, organismal development and disease processes (1,2), regulation of cognition and behavior (3), and personal variation (4). A set of recent efforts established the proof of principle that comparative analyses of interspecies epigenomes can lead to functional annotation of non-coding regulatory genomic sequences (3,5). These discoveries initiated a cohort of exciting studies revealing the evolutionary properties of the mammalian epigenomes and inspired the analyses of co-evolutionary relationships among genomes and epigenomes (5–16). Recently, the comparative studies have been expanded to include chromatin interactomes, providing insights on the evolution of genome organizations and long-range regulatory interactions (17–19). These efforts highlighted how comparative epigenomics, an emerging field, leverages evolutionary patterns of epigenomes to functionally annotate genomes. The rapid growth of functional genomic data also provided ample resources for comparative studies. To date, major epigenome consortia including the RoadMap Epigenomics Project and the ENCODE Project have produced thousands of high-throughput functional genomic datasets in more than 80 tissue types of multiple organisms. Given the advances in analytical methods and the explosive growth of sequencing data, a computational tool for the integrative comparison of the genome and the epigenome is high in demand.

Here we present the EpiAlignment web server, a pairwise alignment tool for both the genome and the epigenome. By providing a pair of epigenomic data, users can align any genomic regions from two species, namely query regions and target regions, with sequences and epigenomic information. The web server incorporates a database of pairwise ChIP-Seq datasets for users’ convenience, which contains peak files of 56 human and 70 mouse ChIP-Seq experiments from 15 matched tissue types and cell lines. Users can either upload their own data or select pre-compiled datasets as inputs for EpiAlignment analyses. EpiAlignment supports two alignment modes: the one-vs-one mode (default) and the many-vs-many mode, corresponding to two types of analyses. By default, the user can align each query region against its corresponding target region to identify a best-matching sub-region with both sequence and epigenomic similarities (Figure 1A). Alternatively, the many-vs-many mode can be used to align each query region against all target regions to find the best match (Supplementary Figure 1). Each mode provides various ways of defining target regions so that the user can easily search against homologous regions or specific gene families for functional comparison (Figure 1B). The alignment results are reported in graphical and tabular formats, with which the user can view alignment scores and best matches found by EpiAlignment. The aligned regions can be further visualized in the UCSC genome browser (20) or the Genomic Interaction Visualization Engine (GIVE) (15) by following links provided in the result tables. With EpiAlignment, users can search for genomic regions that may share similar functions and interrogate functional correspondence among cis-regulatory elements in various tissues between two species.

**Figure 1.**
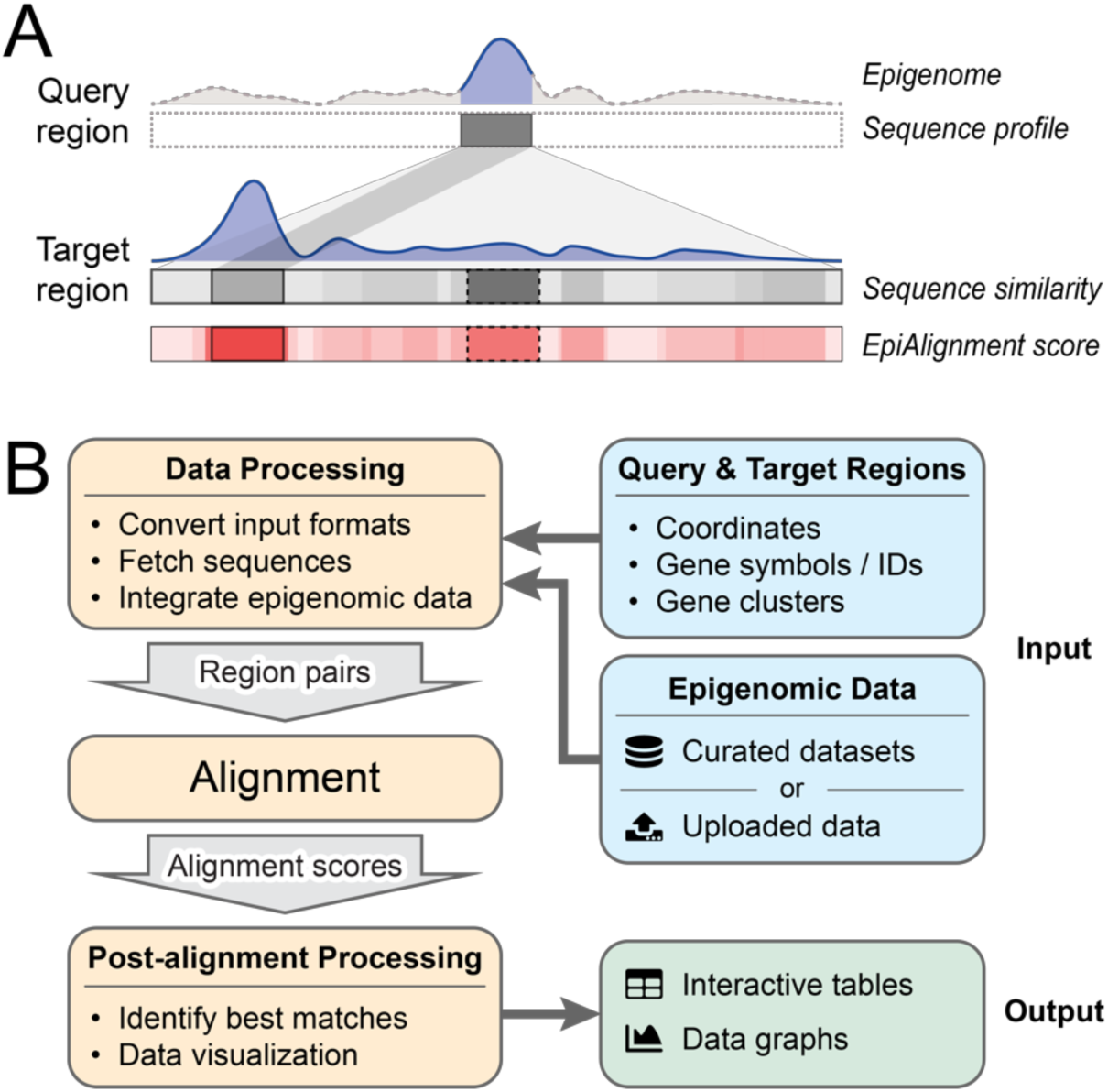
Overview of EpiAlignment. (A) Demonstration of the default alignment mode. The query region (up, dark gray box) and the target region (down, gray bar) are provided by the user as inputs, together with the epigenomic data (purple). The sequence similarity to the query region varies along the target region (grayscale in the “sequence similarity” bar). Within the target region, the gray box with dotted border represents the best sequence match with the highest sequence similarity, and the gray box with solid border represents a sub-region with moderate sequence similarity. In the analysis, EpiAlignment yields scores accounting for both sequence and epigenomic similarities (shades of red in the bottom bar), revealing the best match (red box with solid border) with structural similarity over the best sequence match (red box with dotted border). (B) The EpiAlignment workflow.

## METHODS AND IMPLEMENTATION

### EpiAlignment algorithm

The EpiAlignment algorithm aims to identify the optimal alignment between two genomic regions, denoted as *A* and *B*. Each genomic genome contains two types of data, namely the genomic sequence and the epigenomic profile, denoted as *A_g_* and *A_e_*, respectively. Thus *A* = (*A*_*g*_,*A*_*e*_), and *B* = (*B*_*g*_,*B*_*e*_). We denote each alignment between the two genomic regions as *α*. The following target function is employed to evaluate the quality of alignment *α*:

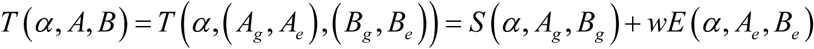

where *T*(*α,A,B*) is the final EpiAlignment score, *S*(*α,A_g_,B_g_*) is the sequence alignment score of the two sequences, *E* (*α*, *A_e_*, *B_e_*) is the similarity score of the two epigenomic profiles, and *w* is the relative weight. When *w* is set to 0, the algorithm degenerates into a sequence-only alignment algorithm. We note that when *α* is integrated out, 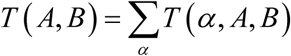corresponds to the log likelihood of *A* and *B* under a recently established genome-epigenome co-evolution model (16). We used the optimization algorithm in (16) to identify the optimal alignment as 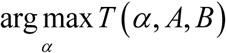.

### Architecture

EpiAlignment is a RESTful service with a tiered structure. A load-balancer powered by Nginx is set up at the front of the service to route requests from multiple users to an appropriate state-less node.js handler. The handler can then spawn up to 140 worker processes and automatically allocate region pairs among them to achieve parallel computing for large datasets from users. The tiers in the architecture are stateless and coupled loosely, allowing easy scaling to better suit variations in user demands.

### Database for pairwise ChIP-Seq experiments

The EpiAlignment web server incorporates a database of pairwise ChIP-Seq experiments, which contains ChIP-Seq datasets curated from the RoadMap Epigenomics Project (21), the ENCODE Project (22,23), the mouse ENCODE Project (5) and published epigenomic comparative studies (24). The current release of EpiAlignment includes 56 human datasets and 70 mouse datasets, obtained from 3 cell lines, 8 adult tissue types and 4 embryonic tissue types (Table 1). Human and mouse tissue types were matched by their bio-sample meta-data, including tissue/organ type, life stage and donor status.

**Table 1.**
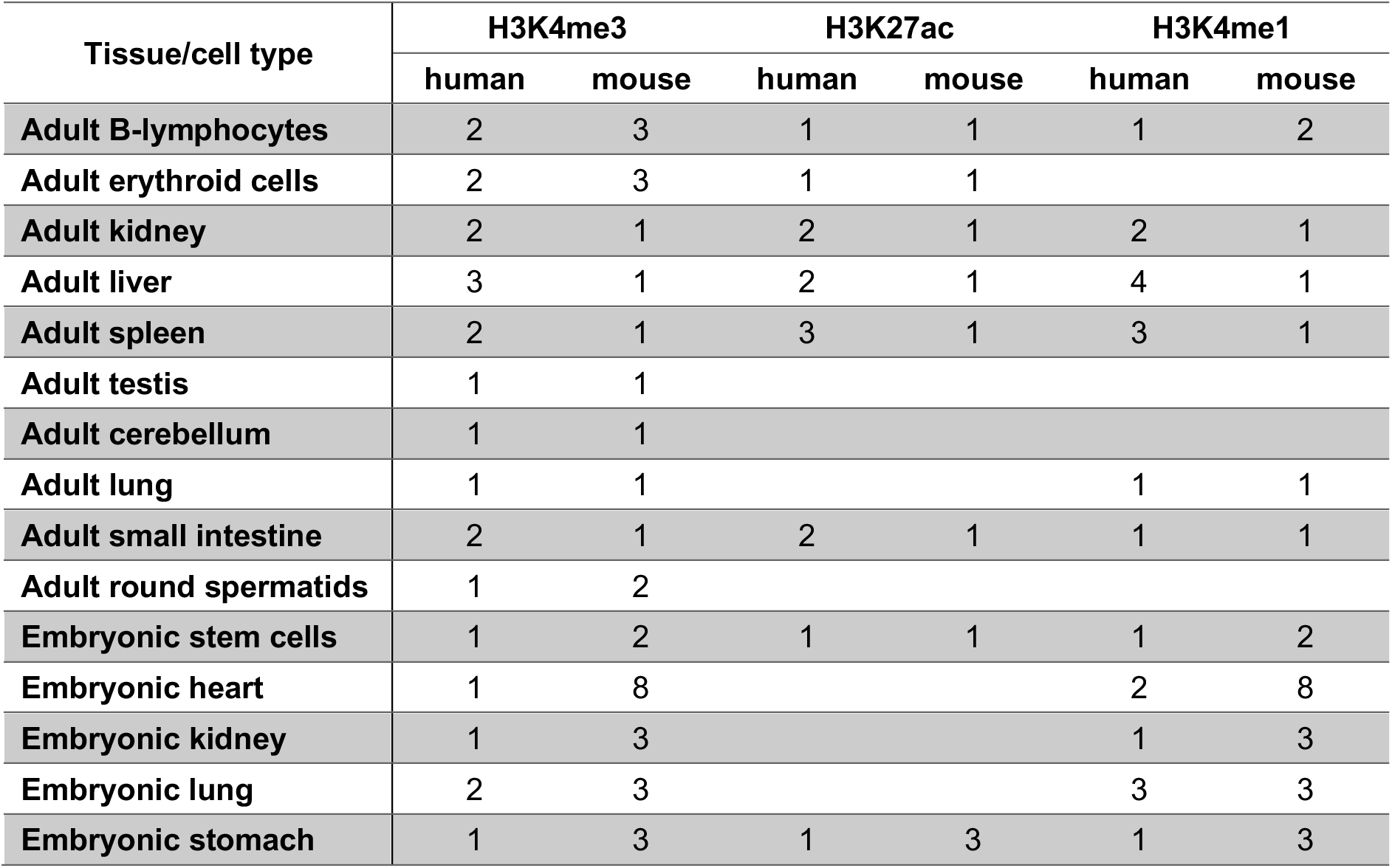
Number of ChIP-Seq datasets in the database for pairwise ChIP-Seq experiments. Rows: cell types or tissues of different life stages. Columns: datasets of different histone modifications, separated by species. Empty cells in the table represent absence of comparable datasets in the corresponding tissue / cell types.

| Tissue/cell type | H3K4me3 |  | H3K27ac |  | H3K4me1 |  |
| --- | --- | --- | --- | --- | --- | --- |
|  | human | mouse | human | mouse | human | mouse |
| Adult B-lymphocytes | 2 | 3 | 1 | 1 | 1 | 2 |
| Adult erythroid cells | 2 | 3 | 1 | 1 |  |  |
| Adult kidney | 2 | 1 | 2 | 1 | 2 | 1 |
| Adult liver | 3 | 1 | 2 | 1 | 4 | 1 |
| Adult spleen | 2 | 1 | 3 | 1 | 3 | 1 |
| Adult testis | 1 | 1 |  |  |  |  |
| Adult cerebellum | 1 | 1 |  |  |  |  |
| Adult lung | 1 | 1 |  |  | 1 | 1 |
| Adult small intestine | 2 | 1 | 2 | 1 | 1 | 1 |
| Adult round spermatids | 1 | 2 |  |  |  |  |
| Embryonic stem cells | 1 | 2 | 1 | 1 | 1 | 2 |
| Embryonic heart | 1 | 8 |  |  | 2 | 8 |
| Embryonic kidney | 1 | 3 |  |  | 1 | 3 |
| Embryonic lung | 2 | 3 |  |  | 3 | 3 |
| Embryonic stomach | 1 | 3 | 1 | 3 | 1 | 3 |

ChIP-Seq experiments for the H3K4me3, H3K27ac and H3K4me1 were selected in the current release because of their prominent role in enhancer and promoter activities. For each ChIP-Seq experiment, only the results containing stable pooled peak regions from multiple replicates and passing the quality control criteria of the consortia were included in the final pairwise database. We also created the metadata on our server to facilitate easy data selection.

### Database for evolutionarily-related genes

EpiAlignment provides a database of evolutionarily-related genes, namely gene clusters, to assist users in functional comparison. The gene clusters were identified by grouping paralogous genes in each species and linking orthologous genes across species. The database contains a total of 2,607 pre-identified gene clusters, including 8,000 human genes and 10,211 mouse genes. Genes without annotated paralogues are not included in the database. The gene clusters are indexed by the gene names and aliases obtained from NCBI annotation database. When users provide a partial name or an Ensembl ID of the gene, EpiAlignment will list all clusters containing the gene(s) with a matching gene name, alias or Ensembl ID for users to choose from as their EpiAlignment targets.

## WEB SERVER DESCRIPTION

### Overview

EpiAlignment provides two major alignment modes: the one-vs-one mode (default) and the many-vs-many mode. The one-vs-one mode attempts to find a best-matching part for a query region in its corresponding target region (one-against-one alignment). It is useful in cases where target regions are much longer than query regions, and best matches of the query regions cannot be easily determined in the target species with sequence homologies only. The many-vs-many mode, on the other hand, was designed to identify the best match for a query region among a set of target regions (one-against-all alignment). It is more useful when the query region corresponds to multiple candidates with potentially similar functions, such as promoters of paralogous genes.

EpiAlignment provides various ways of defining target regions for user convenience. Beside using genomic coordinates directly, users may also take advantage of the specialized methods in each mode to determine the target regions for similarity search. In the default mode, query regions may be mapped against the neighborhood of their homologous regions. In the many-vs-many mode, users may use gene promoters as input regions and map query promoters against promoters of a group of evolutionarily-related genes, namely a gene cluster, for functional comparison. An overview of the alignment modes and supported target region types is shown in Table 2.

**Table 2.**
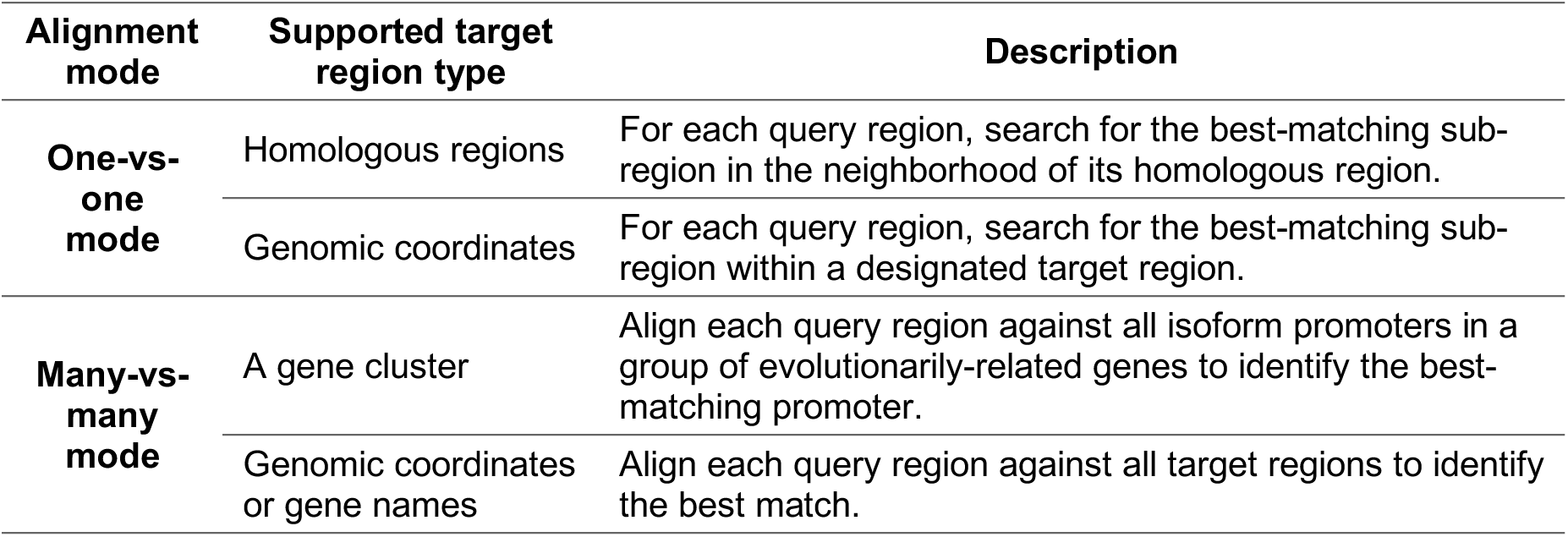
Alignment modes and supported target region types.

### Input

EpiAlignment uses the one-vs-one mode by default, whereas the user may switch to the many-vs-many mode before starting an EpiAlignment run. In both modes, EpiAlignment requires two types of inputs from users: (1) a pair of peak files from ChIP-Seq experiments, and (2) query and target regions to be aligned, both from two species. With the genome assemblies specified, users can either upload their own peak files or select a pair of epigenomic datasets from the database of pairwise ChIP-Seq experiments. Details of each ChIP-Seq experiment can be viewed on the ENCODE or the GEO website by following the links provided in the database. The query and target regions can be provided in various ways depending on the alignment mode used.

#### Specifying query and target regions in one-vs-one mode

In the default mode, the query and target regions should be provided as two lists of genomic coordinates in BED6 format, in which the corresponding lines specify query-target region pairs. The coordinates can either be pasted in the text area of the “Query regions” and “Target regions” boxes or uploaded as files. The web server also allows users to define target regions based on homologous information. In this case, users only need to specify the number of base pairs by which the query regions will be increased. The web server will extend each query region and remap it to the target species to define the corresponding target region.

#### Specifying query and target regions in many-vs-many mode

In this mode, the query regions can be provided as genomic coordinates in BED6 format. Gene symbols or Ensembl IDs are also accepted if gene promoters are used as input regions. When using gene symbols or Ensembl IDs, ranges of promoters need to be specified and the web server will fetch promoter sequences automatically. By default, the (−1,000, +500) flanking regions around TSSs are used as promoter regions. The target regions can be specified in the same way in the “Target regions” box or by selecting a preset gene cluster. Users can search for a gene cluster by typing a gene symbol or an Ensembl ID in the searching bar. All gene promoters in the cluster will be used as target regions.

With the query and target regions defined, the user may adjust the epigenome weight and alignment parameters. This step is optional as default parameters are provided on the webpage. Finally, the user can initiate the alignment by clicking the “Submit” button.

### Output

The user will be redirected to the result page after successfully submitting the data. The result page will refresh automatically until the task is done. An URL to the page will also be sent to the user if an email address was provided.

In the default mode, an interactive table will be returned, in which each row corresponds to one query-target region pair. The user may click on the row to expand it and view the alignment score distributions along the target region. In both modes, all genomic regions can be viewed individually in the UCSC genome browser by clicking the icon beside each genomic coordinate in the tables, or in region pairs in the Genomic Interaction Visualization Engine (GIVE) by clicking the icon at the beginning of the rows. The user may select ChIP-Seq data from the custom tracks and zoom/slide the view window of the browser to further investigate the genomic contexts of the regions.

In the many-vs-many mode, two heatmaps and a table with detailed results will be returned. The heatmaps illustrate alignment scores between query regions (rows) and target regions (columns), yielded with and without epigenomic information. For each query region, EpiAlignment detects if its best-matching target region switched from one to another between alignments with and without the epigenome. If the hit has altered, the best hits found with and without epigenomic information are highlighted by green and gray boxes respectively. Details of the alignments are listed in the table on the same page, which can be downloaded by clicking “Download results”.

### Runtime and computational performance

The time complexity of each alignment is O(*mn*) where *m* and *n* are the lengths of the sequences to be aligned. The architecture of EpiAlignment utilizes parallel computing of up to 140 worker threads. The runtime for a typical input set in the default mode (50 region pairs, about 2,000bp for query regions and 40,000bp for target regions) is about 2 minutes, and the runtime for a typical input set in the many-vs-many mode (50 region pairs, about 1500bp for both query and target regions) is under 15 seconds.

## DATA APPLICATIONS

Two examples with sample inputs were provided on the EpiAlignment website intended for users to get familiar with the two alignment modes.

### Case study 1

This example demonstrated how to find regions with similar chromosomal structures using the default one-vs-one mode. The promoter regions of human gene *LDAH* and its orthologous mouse gene *Ldah* are both marked by H3K4me3 in round spermatids, whereas the underlying sequences are poorly conserved. With liftOver (25), a sequence homology-based tool, the mouse promoter was mapped to a human region approximately 10 kb from human *LDAH* promoter, and the result human region showed no H3K4me3 occupancy. We used the H3K4me3-marked region in mouse *Ldah*’s promoter as the input query region (chr12:8,207,583-8,209,349) and aligned it against the −20,000 ~ +20,000 neighborhood of its homologous region in human. The H3K4me3 ChIP-Seq datasets of human and mouse round spermatids were selected from the database as epigenomic information (24). The sequence-only alignment (*w*=0) revealed two hits with similar alignment scores. The downstream one corresponded to *LDAH*’s promoter, while it showed an alignment score slightly lower than the best sequence match. With the H3K4me3 information, the best hit shifted to *LDAH*’s promoter, and EpiAlignment successfully matched the two promoter regions together (Figure 2). Given the orthologous relationship between the two genes, it is more reasonable that their promoters share similar functions. This case illustrates how EpiAlignment can be used to identify functional genomic sites with structural similarities and infer functional correspondence in regions lacking sequence conservation.

**Figure 2.**
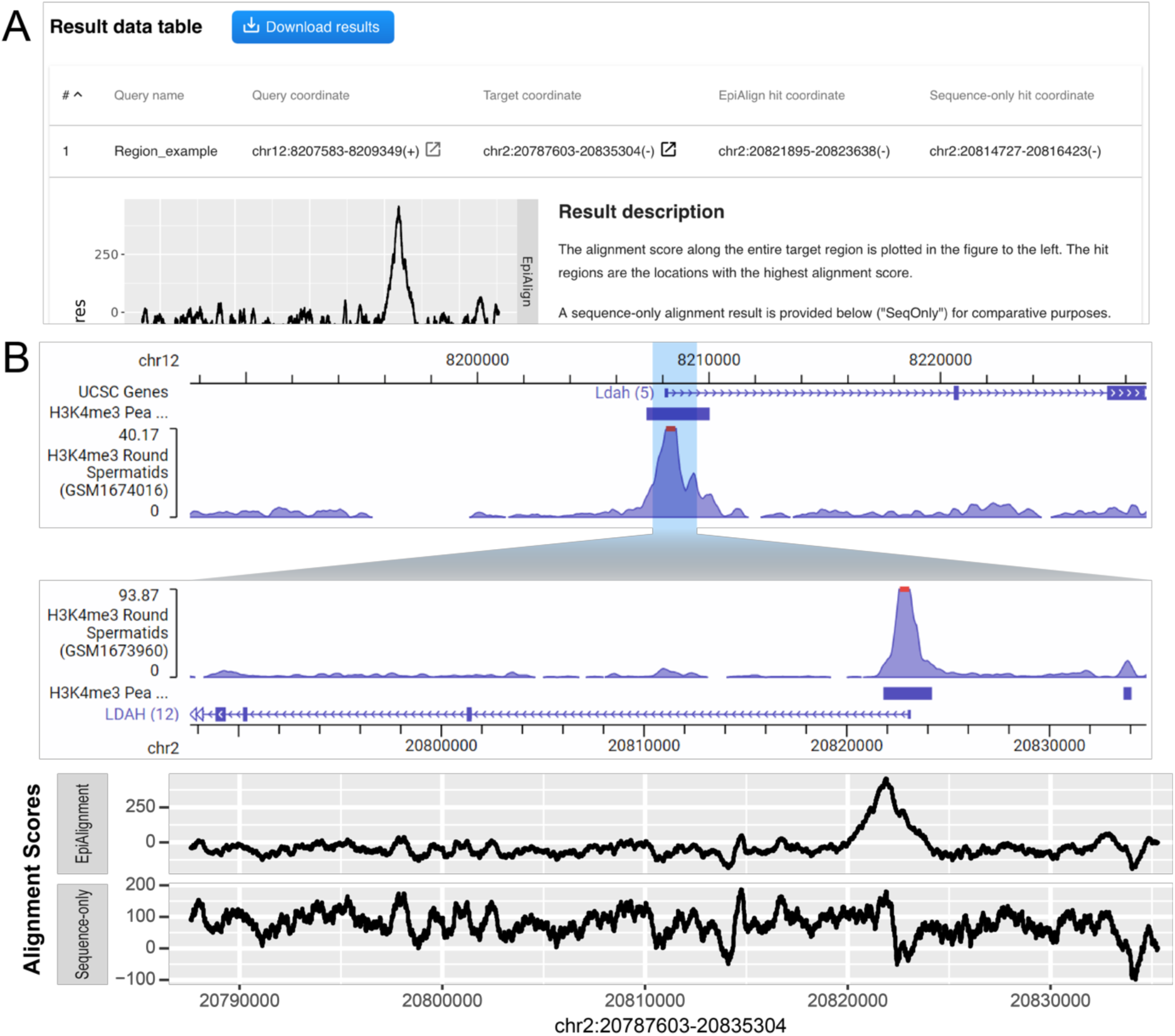
One-vs-one mode results. (A) The interactive table containing coordinates of input regions and best matches. The user can click on each row to expand it and view detailed results. The user may also view the query and target regions together in GIVE by clicking the icon at the beginning of each row (solid red box), or individually by clicking the icon behind each coordinate (dotted red box). (B) Visualization of the input query and target regions in GIVE, together with alignment scores along the target region. Upper: screenshot of GIVE showing the query (blue-shaded area at top) and target (bottom) regions. Lower: distribution of alignment scores along the target region, produced with (top) and without (bottom) epigenomic information. X-axis: genomic coordinates aligned to the GIVE view. Y-axis: alignment scores.

### Case study 2

In this example, we aligned the promoter of a human protocadherin beta gene, *PCDHB5*, against a mouse gene cluster using the many-vs-many mode. The same epigenomic datasets were used as in Case study 1. We used the gene symbol *PCDHB5* as input to define the query region. In the “Target regions” box, we selected “Search a gene cluster”, searched PCDHB5 and chose “Cluster_1491” in the pop-up panel as input. The cluster contains six mouse protocadherin beta genes: *Pcdhb4*, *Pcdhb6*, *Pcdhb8*, *Pcdhb10, Pcdhb11* and *Pcdhb12*, all of which are annotated orthologous genes of *PCDHB5*. In sequence-only alignment (*w*=0), the promoter of *Pcdhb11* exhibited the highest sequence similarity to *PCDHB5*’s promoter. After incorporating the H3K4me3 data, the best hit switched to *Pcdhb12*’s promoter (Figure 3A, B). The alignment score between *PCDHB5* and *Pcdhb12*’s promoters increased after taking the epigenome into account, suggesting that the two promoters had consistent H3K4me3 occupancies. The visualization of results showed that both *PCDHB5*’s and *Pcdhb12*’s promoters exhibited significant H3K4me3 signals, whereas *Pcdhb11*’s promoter harbored no H3K4me3 peak (Figure 3C). This example illustrates that by finding structural similarities among promoters, EpiAlignment may help to shed light on potential functional correspondence among evolutionarily-related genes.

**Figure 3.**
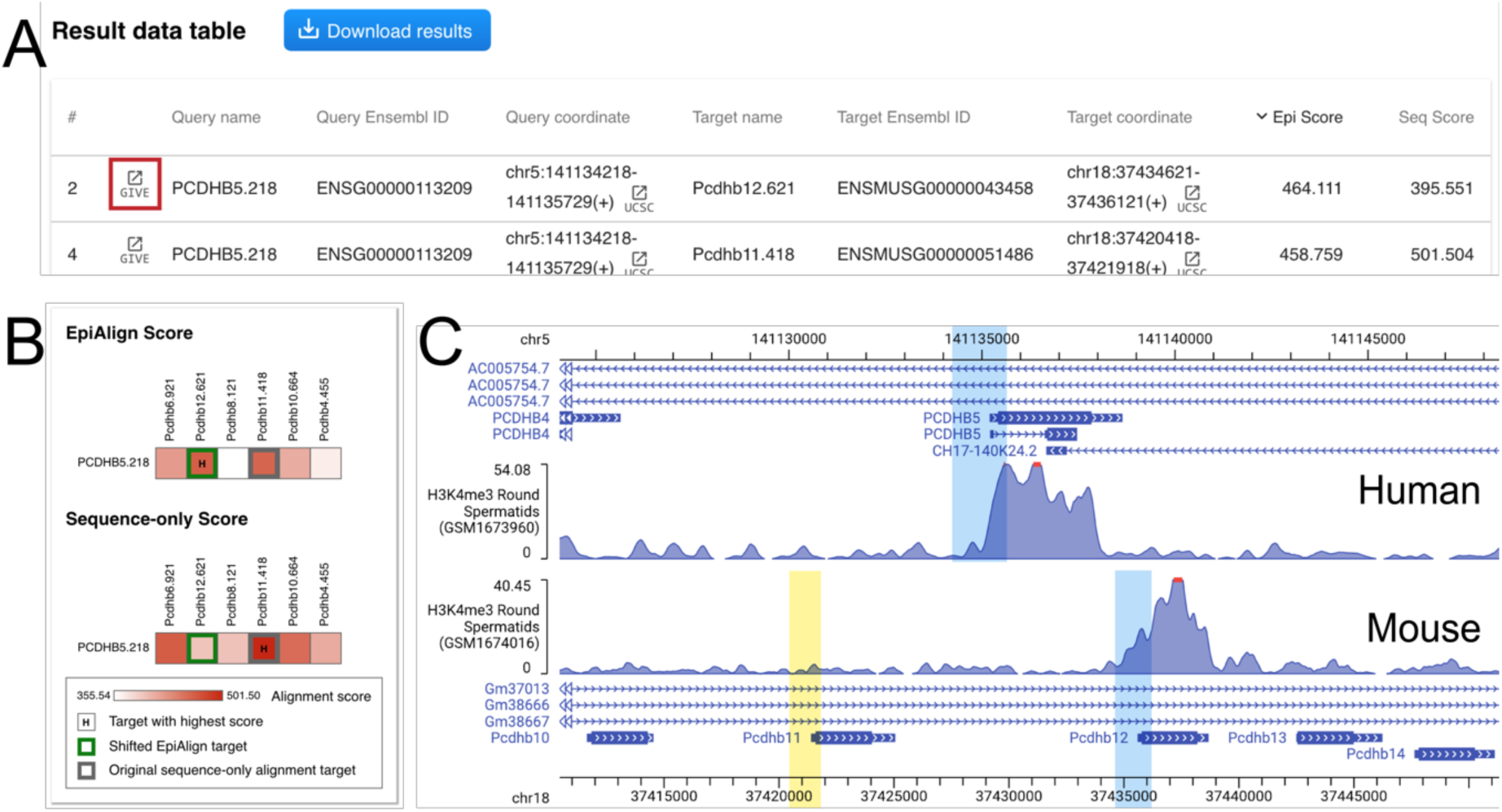
Many-vs-many mode results. (A) The interactive table containing coordinates of input regions and best matches, alignment scores and links for visualization. The user may sort the table by each column by clicking on the column name. (B) Heatmaps of alignment scores on the result page. Rows: query regions. Columns: target regions. Color-scale in the cells: alignment scores of query-target region pairs. Green box: the best match found with epigenomic information. Gray box: the best match found with sequence only. “H” in the cell: the target region with the highest alignment score in the row. (C) Visualization of the regions in GIVE. Top: the genomic context of the input query region (blue). Bottom: the genomic context of the best matches found by EpiAlignment, with (blue) and without (yellow) epigenomic information.

## DISCUSSION

Integrative analyses of the genome and the epigenome can provide insights for the annotation of regulatory sequences, especially in regions lacking sequence conservation. We have developed EpiAlignment, a cross-species alignment tool incorporating both the genome and the epigenome. By assessing structural similarities between functional genomic sites, the tool may assist researchers in identifying functional correspondence across species, which provides a starting point for downstream experiments and analyses. The interface of EpiAlignment is designed to be user-friendly and does not require users to be experienced in computational biology. It also shows versatility in providing various modes and supporting different input types. Users may easily use the one-vs-one mode on homologous neighborhoods of regulatory elements to refine their annotations, or search for a pre-defined gene cluster and align them with the many-vs-many mode to study the potential functional correspondence among evolutionarily-related genes.

Given that EpiAlignment uses peak files from epigenomic experiments to assess epigenomic signatures within regions, its output can be affected by the peak-calling results. Therefore, we only included stable peak files from ENCODE experiments with replicates pooled in our database for pairwise ChIP-Seq experiments. When using custom data, users may adjust thresholds for peak-calling and examine the genomic regions surrounding the peaks in genome browsers for better performance.

EpiAlignment can be further enhanced in several aspects in the future. First, the database for pairwise ChIP-Seq experiments can be expanded to include more comparable datasets of different experiment types, including DNase-Seq, ATAC-Seq and MeDIP-Seq. Species other than human or mouse can also be supported in future updates when sufficient epigenomic data is available. Further expansion of the algorithm may allow EpiAlignment to align with multiple epigenomic marks simultaneously to better reflect the interactions among different epigenomic modifications. Moreover, while the current release of EpiAlignment does not support genome-wide alignments due to speed constraints, future investments in computational hardware and algorithm developments may improve the speed performance and enable alignments against the entire genome.

## SUPPLEMENTARY DATA

Supplementary Data are available at NAR online.

## ACKNOWLEDGEMENT

We thank Dr. Zhangming Yan for useful discussions.

