## Supplementary Figure 1 for "EpiAlignment: alignment with both DNA sequence and epigenomic data"

**SUPPLEMENTARY DATA**

**Supplementary Figure 1. Demonstration of the many-vs-many mode.** Left: query and target regions are provided as inputs by the user, together with the epigenomic data (purple). Sequence similarities vary among the query and target regions (grayscale in bars). Each query region is aligned against all target regions (dashed double-headed arrows). Right: The EpiAlignment output. The alignment score is shown in red color for each pair of query region (row) and target region (column).

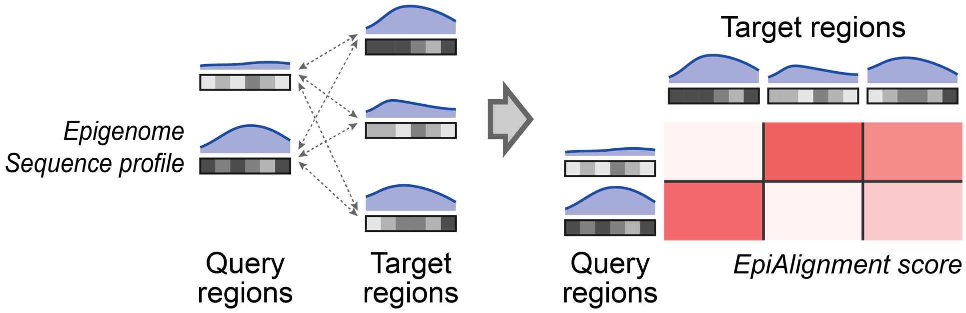
